# A *Drosophila* Insulator Interacting Protein Suppresses Enhancer-Blocking Function and Modulates Replication Timing

**DOI:** 10.1101/661041

**Authors:** Emily C. Stow, Ran An, Todd A. Schoborg, Nastasya M. Davenport, James R. Simmons, Mariano Labrador

## Abstract

Insulators play important roles in genome structure and function in *Drosophila* and mammals. More than six different insulator proteins are required in *Drosophila* for normal genome function, whereas CTCF is the only identified protein contributing to insulator function in mammals. Interactions between a DNA binding insulator protein and its interacting partner proteins define the properties of each insulator site. The different roles of insulator protein partners in the *Drosophila* genome and how they confer functional specificity remain poorly understood. Functional analysis of insulator partner proteins in *Drosophila* is necessary to understand how genomes are compartmentalized and the roles that different insulators play in genome function. In *Drosophila*, the Suppressor of Hairy wing [Su(Hw)] insulator is targeted to the nuclear lamina, preferentially localizes at euchromatin/heterochromatin boundaries, and is associated with the *Gypsy* retrotransposon. The properties that the insulator confers to these sites rely on the ability of the Su(Hw) protein to bind the DNA at specific sites and interact with Mod(mdg4)-67.2 and CP190 partner proteins. HP1 and insulator partner protein 1 (HIPP1) is a recently identified partner of Su(Hw), but how HIPP1 contributes to the function of Su(Hw) insulators has not yet been elucidated. Here, we find that mutations in the HIPP1 crotonase-like domain have no impact on the function of Su(Hw) enhancer-blocking activity but do exhibit an impaired ability to repair double-strand breaks. Additionally, we find that the overexpression of each HIPP1 and Su(Hw) causes defects in cell proliferation by limiting the progression of DNA replication. We also find that HIPP1 overexpression suppresses the Su(Hw) insulator enhancer-blocking function.

## Introduction

Chromatin insulator proteins function by coordinating the regulation of gene expression with chromosome structure. Canonical roles of insulator proteins include their ability to prevent communication between enhancers and target promoters and their role in forming boundaries between regions of heterochromatin and euchromatin. (Wallace and Felsenfeld, 2007; West *et al*., 2002; Yang and Corces, 2012). The *Drosophila melanogaster* genome encodes a diverse array of insulator proteins, each with unique roles and binding sites contributing to both genome structure and transcriptional regulation. Accessory proteins that interact with DNA binding insulator proteins are essential for insulator function, and the discovery of novel insulator partner proteins may contribute to our understanding of insulator function and mechanisms.

The 5’ untranslated region of the *Gypsy* retrotransposon contains 12 binding sites for the Suppressor of Hairy wing [Su(Hw)] protein, allowing *Gypsy* to function as an insulator in *Gypsy* insertion sites. Flies with mutations in *su(Hw)* have no discernable phenotype other than female sterility due to oogenesis-specific phenotypes (Hsu *et al*., 2019; Hsu *et al*., 2015; Klug *et al*., 1968; Klug *et al*., 1970; Soshnev *et al*., 2013). All insulator binding proteins rely on partner proteins to carry out basic insulator functions. Thus, Centrosomal Protein 190 (CP190) and Modifier of mdg4 67.2 [Mod(mdg4)] are Broad-complex, Tramtrack and Bric-a-brac/Poxvirus, and Zinc Finger (BTB/POZ) domain-containing proteins that interact with Su(Hw) to form the Su(Hw) insulator complex (Georgiev and Gerasimova, 1989; Gerasimova *et al*., 1995; Pai *et al*., 2004). Both Mod(mdg4) and CP190 are required for the enhancer blocking function of the Su(Hw) insulator (Gerasimova *et al*., 1995; Kurshakova *et al*., 2007; Pai *et al*., 2004).

A recent study identified yet another member of the *Gypsy* insulator complex called Heterochromatin protein 1 and Insulator Partner Protein 1 (HIPP1) (Alekseyenko *et al*., 2014). HIPP1 is found to interact with multiple DNA-binding protein complexes, including a high-confidence association with Heterochromatin Protein 1a (HP1) and insulator proteins CTCF and Su(Hw). Another recent study investigated the localization and developmental patterns of HIPP1 (Glenn and Geyer, 2018). This study found that HIPP1 is primarily recruited to euchromatin regions in a Su(Hw)-dependent manner and that *hipp1* null mutants have no discernable phenotype and that mutation of *hipp1* does not affect the function of the Su(Hw) insulator. This study also identified HIPP1 as the fly homolog of the human Chromodomain Y-like protein (CDYL). Both HIPP1 and CDYL contain a crotonase-like domain (CLD), which is able to mediate interactions with histone modifiers to prevent acetylation and crotonylation while promoting the addition of repressive histone modifications (Wu *et al*., 2009). CDYL exists in multiple isoforms. The CDYLb isoform contains an N-terminal chromodomain in addition to its C-terminal CLD. HIPP1 does not contain a chromodomain and therefore shares more similarities with the CDYLa and CDYLc isoforms, which also lack a functional N-terminal chromodomain (Glenn and Geyer, 2018; Wu *et al*., 2013). The human CDYL protein has been found to associate with histone remodeling complexes to promote heterochromatin formation and maintenance, including interactions with the Polycomb Repressive Complex 2 (PRC2) and chromatin assembly factor-1 (CAF-1) (Liu *et al*., 2017b; Zhang *et al*., 2011). CDYL specifically associates with H3K9me3 as well as di- and tri-methylated H3K27 and negatively regulates lysine crotonylation, a modification associated with active promoters (Franz *et al*., 2009; Liu *et al*., 2017a; Zhang *et al*., 2011). CDYL is also a component of the Repressor element-1 silencing transcription factor (REST) complex. CDYL contributes to REST-mediated transcriptional silencing of target genes by mediating the interaction between REST and the H3K9-specific G9a methyltransferase (Zhang *et al*., 2011).

In addition to its insulator activity, Su(Hw) appears to function as a transcriptional regulator of neural genes and has previously been suggested as a functional homolog of mammalian REST (Lakowski *et al*., 2006; Soshnev *et al*., 2012). Additionally, the sterility phenotype of *su(Hw)* mutant females has been linked to the de-repression of neural genes in the female germline (Soshnev *et al*., 2013). The association of HIPP1, a CLD-containing protein, with Su(Hw) adds to the similarities between Su(Hw) and the mammalian REST complex, however a recent study (Glenn and Geyer, 2013) did not find a significant effect on the expression of Su(Hw)-regulated genes in HIPP1 mutants. This suggests that the CLD function of HIPP1 may contribute to alternative roles of the Su(Hw) complex.

One recent study demonstrates that CDYL localizes to sites of double strand breaks (DSBs) to promote recruitment of the Polycomb Repressive Complex 2 subunit Enhancer of Zeste 2 (EZH2), leading to transcriptional repression through trimethylation of H3K27 (Abu-Zhayia *et al*., 2018). This study further reveals that CDYL recruitment to DSBs occurs normally in mutants for the chromodomain and concludes that this role for the CDYL protein is dependent on the CLD. Additionally, CDYL-depleted cells exhibit a reduced accumulation of H3K27me3 at DSB sites as well as a heightened sensitivity to DNA damage-inducing agents such as ionizing radiation and the chemotherapy drug cisplatin. CDYL-depleted cells also exhibit a significant reduction in homology directed repair (HDR) frequency, with no significant change in the frequency of break sites repaired by non-homologous end joining (NHEJ). Since the *Drosophila* HIPP1 protein contains a CLD homologous to the CLD of human CDYL and lacks a functional chromodomain, it is possible that the HIPP1 protein plays a role similar to the CDYL protein in transcriptional silencing at DSBs and in promotion of the HDR pathway (Abu-Zhayia *et al*., 2018).

Chromatin insulators are important components of genome architecture across Eukaryotes (Chung *et al*., 1997; Farrell *et al*., 2002; Guo *et al*., 2008; Heger *et al*., 2013; Heger *et al*., 2012; Heger *et al*., 2009; Hily *et al*., 2009; Ishii *et al*., 2002; Ishii and Laemmli, 2003; Palla *et al*., 1997; Schoborg and Labrador, 2010). Recent advances in mammalian systems have pinpointed roles for the insulator protein CTCF in shaping genome architecture in a flexible manner to allow for changes in gene transcription and dynamics of the DNA fiber as the cell cycle progresses (Belton *et al*., 2012; Fudenberg *et al*., 2016; Naumova *et al*., 2013). The loop extrusion model involves the extrusion of DNA loops by the cohesin complex until the complex encounters and forms a stable interaction with two CTCF molecules bound to DNA in opposing orientation. The formation of stable loops creates topologically associating domains (TADs) with CTCF sites located at the border between consecutive TADs, in which the CTCF-cohesin loop anchor colocalizes with break point cluster regions (BCRs) generated by Topoisomerase 2B (TOP2B) (Canela *et al*., 2017). BCRs at loop anchors are thought to occur due to the torsional strain accumulated during transcription, replication, and folding of the genome. The colocalization between loop anchors and BCRs suggests loop extrusion mediated by cohesin and CTCF places a conformational strain on the nucleus that must be alleviated by TOP2B activity.

Although the loop extrusion process has not yet been described in *Drosophila*, similarities between mammalian CTCF and *Drosophila* insulator proteins encourage investigation into conserved mechanisms. It has been shown that function and stabilization of the Su(Hw) insulator complex relies on Topoisomerase 2 (TOP2) activity, specifically through an interaction between TOP2 and Mod(mdg4)2.2 that stabilizes the association of Mod(mdg4)2.2 with the Su(Hw) complex (Ramos *et al*., 2011). Additionally, Su(Hw) interacts with *Drosophila* Topoisomerase I-Interacting Protein (dTopors), a protein that binds the nuclear lamina and directs Su(Hw) binding sites to positions along the nuclear lamina in order to define lamina associated domains (LADs) (Capelson and Corces, 2005). The association of genomic sites with the nuclear lamina confines movement of the DNA fiber during processes such as transcription and replication (Gonzalez-Sandoval and Gasser, 2016). It has not yet been shown how Su(Hw) binding is regulated to allow transcription or replication of Su(Hw)-bound sequences located in LADs.

In agreement with the well-documented association of Su(Hw) with band and interband transitions, it has been shown that Su(Hw) binding sites are enriched in malachite chromosome fragments which are regions flanking intercalary heterochromatin (IH) domains (Khoroshko *et al*., 2016). IH domains resemble pericentric heterochromatin but are found interspersed in the euchromatic regions of the genome. These domains were originally identified as the bands along arms of polytene chromosomes from *Drosophila* salivary glands (Belyaeva *et al*., 2008; Kaufmann, 1939). Replication of IH domains occurs late during the replication timing program and is initiated by origins in the surrounding euchromatin (Lubelsky *et al*., 2014; Pope *et al*., 2014). The flanking malachite regions containing Su(Hw) replicate first, followed by the internal IH content. The positioning of Su(Hw) binding sites in these transition regions between euchromatin and heterochromatin suggests that the Su(Hw) protein complex may be regulated in a cell cycle-specific manner to allow entry of replication machinery into intercalary heterochromatin. In agreement with this model, we recently reported that mutations in *su(Hw)* contribute to replication stress in developing *Drosophila* egg chambers and dividing neuroblasts (Hsu *et al*., 2019). The mechanism by which Su(Hw) is required to maintain genome stability during DNA replication has not yet been elucidated.

The identification of a novel Su(Hw)-interacting protein such as HIPP1 provides an opportunity to further investigate *Drosophila* insulator mechanisms and functions. Here, we analyze the relationship between HIPP1 and Su(Hw) and provide evidence of novel roles for the Su(Hw) insulator complex in cell proliferation and genome stability. We have developed fly lines overexpressing Su(Hw), lines overexpressing HIPP1, and a CRISPR-generated mutant of endogenous *hipp1* with a deletion of the CLD domain (*hipp1^ΔCLD^)*. We show that Su(Hw) and HIPP1 overexpression result in severe cell proliferation defects. Overexpressing HIPP1 also results in the excess accumulation of larval brain cells in the early phase of DNA replication, suggesting HIPP1 expression levels may regulate phases of the replication timing program in *Drosophila*. We additionally provide evidence that larval brain cells from *hipp1^ΔCLD^* mutants are deficient in DNA damage repair following X-ray irradiation. We also show that overexpression of HIPP1 results in suppression of Su(Hw)-mediated enhancer blocking with no disruption of Su(Hw) binding to DNA. This disruption of insulator function correlates with the displacement of cohesin from the Su(Hw) sites. These results provide additional evidence that Su(Hw) plays a role in cell proliferation that may be dependent on a role in DNA replication. This study also provides compelling evidence that HIPP1 functions in the DNA damage repair pathway in a similar manner as the human CLD-containing protein, CDYL. Taken together, these findings further suggest insulator proteins contribute important functions to the processes of genome replication and genome stability, raising new and intriguing questions about the mechanisms mediating such functions.

## Results

### *hipp1* mutants lacking the crotonase-like domain are deficient in DNA repair

HIPP1 has been identified as the homolog of the human CDYL protein (Glenn and Geyer, 2018). The HIPP1 and CDYL proteins both contain a C-terminal crotonase-like domain (CLD) while CDYL also contains an N-terminal chromodomain. The presence of conserved features between the CDYL and HIPP1 crotonase-like domains suggests that the function of this domain is consistent between the two proteins. Both CDYL and HIPP1 contain critical residues to form an oxyanion hole which is required for stabilizing an anion intermediate produced during reactions with an acyl-CoA substrate (Glenn and Geyer, 2018; Wu *et al*., 2009). Figure 1A shows the alignment of these critical residues, indicated by red boxes, between HIPP1 and CDYLb, the most abundant isoform of CDYL (Abu-Zhayia *et al*., 2018). The function of human CDYL in promoting the HDR pathway occurs normally in chromodomain mutants, suggesting this role relies on the crotonase-like domain. Therefore, it is possible that HIPP1 shares this role with CDYL.

**Figure 1.**
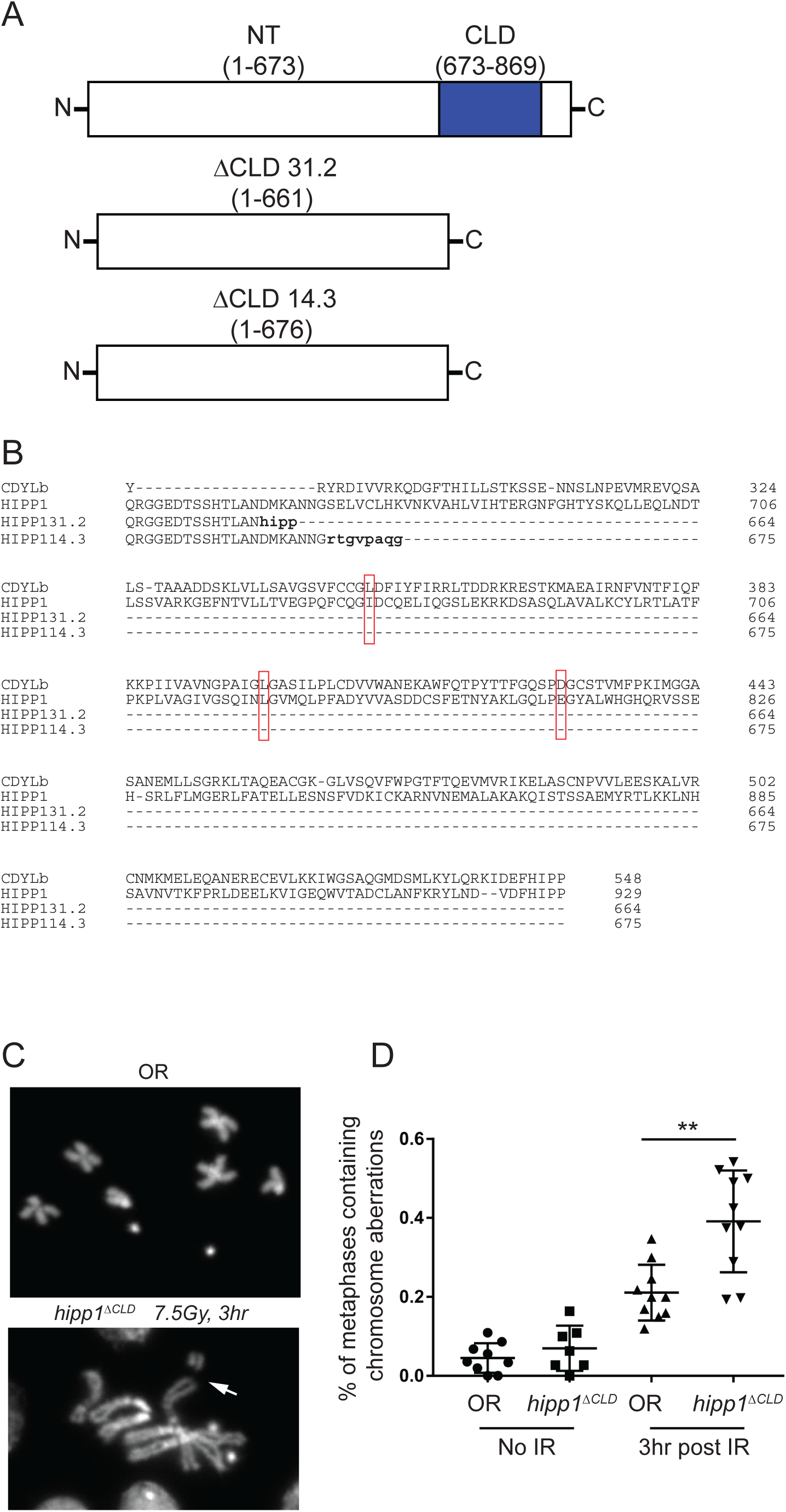
Mutations in HIPP1 disrupt DNA damage repair pathways. (A) A wild-type and two *hipp1* mutant alleles are shown. The crotonase-like domain is shown in blue. Mutant alleles for *hipp1*, 31.2 and 14.3, contain a frameshift (followed by an early stop codon) preceding the crotonase-like domain. (B) A sequence alignment of human CDYLb, wild-type HIPP1, and our *hipp1* mutant alleles are shown. The critical residues for forming the oxyanion hole are outlined in red (L403, L452, D483). (C) Representative images of an Oregon-R chromosome spread (top) and *hipp1^ΔCLD^*chromosome spread (bottom) from *Drosophila* brains are shown. The Oregon-R spread contains no chromosome aberrations (CABs) while the *hipp1^ΔCLD^* chromosomes show one CAB indicated by the white arrow. (D) The percent of metaphases containing one or more CABs for each condition is shown. Both Oregon-R and *hipp1^ΔCLD^* show an increase in CABs following irradiation (IR) and recovery. *hipp1^ΔCLD^*displays a significant increase in the number of CABs, compared to Oregon-R following irradation and recovery (p=0.0017). Statistical Significance was determined using an unpaired t-test.

To investigate whether the CLD of CDYL and HIPP1 play a similar role in the HDR pathway, we generated mutants by specifically removing the CLD domain of HIPP1 using CRISPR-Cas9. We targeted guide RNAs to sequences flanking the region encoding the CLD and generated a deletion and stop codon early in the sequence. We generated two different CLD deletion fly lines (HIPP1 31.2 and HIPP1 14.3, Figure1A). Both fly lines contain a frameshift followed by a stop codon early in the CLD-encoding region and lack the residues critical for the formation of an oxyanion hole (Glenn and Geyer, 2018; Wu *et al*., 2009). The generation of early stop codons and deletion of the CLD in both alleles was confirmed by DNA and cDNA sequencing (data not shown). Figure 1A shows the alignment of our mutant alleles with the CLD from Oregon-R *hipp1* and human CDYLb. We performed experiments using flies *trans*-heterozygous for the HIPP1 31.2 and 14.3 to limit any effect from possible off-target mutations induced by the CRISPR Cas-9 method.

Next, to determine whether HIPP1 participates in the HDR pathway, we evaluated the ability of dividing neuroblasts from *hipp1^ΔCLD^*larval brains to recover from DNA damage by quantifying the occurrence of chromosomal abberations (CABs) following X-ray treatment and recovery (Gatti and Goldberg, 1991; Merigliano *et al*., 2017). CABs were quantified by counting the number of metaphasic nuclei containing one or more aberrations and comparing this number to the total number of metaphases, including those with no CABs (Figure 1B). We found that *hipp1^ΔCLD^* samples contained a significantly higher number of metaphases with one or more CABs following X-ray treatment and recovery compared to the Oregon-R control (Figure 1D, p=0.0017). This result suggests that a role for the CLD in DNA damage repair is conserved between the human CDYL and *Drosophila* HIPP1 proteins. How this role relates to HIPP1-containing complexes remains unknown. There is evidence of a relationship between DNA damage repair and the CTCF insulator protein in humans, but there is limited evidence that Su(Hw) participates in DNA damage repair (Lang *et al*., 2017; Lankenau *et al*., 2000). It will require additional studies to determine a link between this role of HIPP1 and the Su(Hw) insulator complex, however this conserved function further supports the idea that HIPP1 and CDYL are homologous proteins.

### Su(Hw) and HIPP1 colocalize and dynamically bind to polytene chromosomes

Functional analysis of different *su(Hw)* mutations and the genomic distribution of Su(Hw) binding sites suggests that different binding sites may have different functions, depending on the genomic location and the partner proteins associated with Su(Hw) at the given site (Soshnev *et al*., 2012). To further characterize the interaction between Su(Hw) and HIPP1, we developed Gal4-inducible transgenic constructs fused to fluorescent proteins to observe localization patterns of HIPP1 (P{HIPP1::mc, w^+^}) relative to Su(Hw) (P{Su(Hw):: GFP, w^+^}) binding sites (Figure 2A). We drove the expression of the transgenic constructs with a *vestigial* Gal-4 promoter (*vg*-Gal4) that specifically drives transgenic expression in larval wing discs but also induces significant expression in salivary glands (Barwell *et al*., 2017; Schoborg *et al*., 2013). We immunostained with antibodies specific for GFP and RFP (mCherry) to observe the localization patterns of Su(Hw)::GFP and HIPP1::mc on polytene chromosomes from larval salivary glands. Under these conditions, we observed a significant overlap between Su(Hw) and HIPP1 signal (Figure 2B). Next, we analyzed localization patterns of HIPP1::mc and Su(Hw):: GFP in S2 cells in both normal media and osmotic stress media (growth media supplemented with 250mM NaCl). Osmotic pressure drives the formation of insulator bodies in *Drosophila* cells, and all kown *Drosophila* insulator proteins associate with these bodies (Schoborg *et al*., 2013). Therefore, we can indirectly ask whether HIPP1 is associated with insulator function by determining whether HIPP1 also localizes to insulator bodies. We found that Su(Hw)::GFP and HIPP1::mc staining patterns overlap with insulator bodies, following the addition of osmotic stress media, supporting the notion that HIPP1 is closely associated with insulator function (Figure 2C).

**Figure 2.**
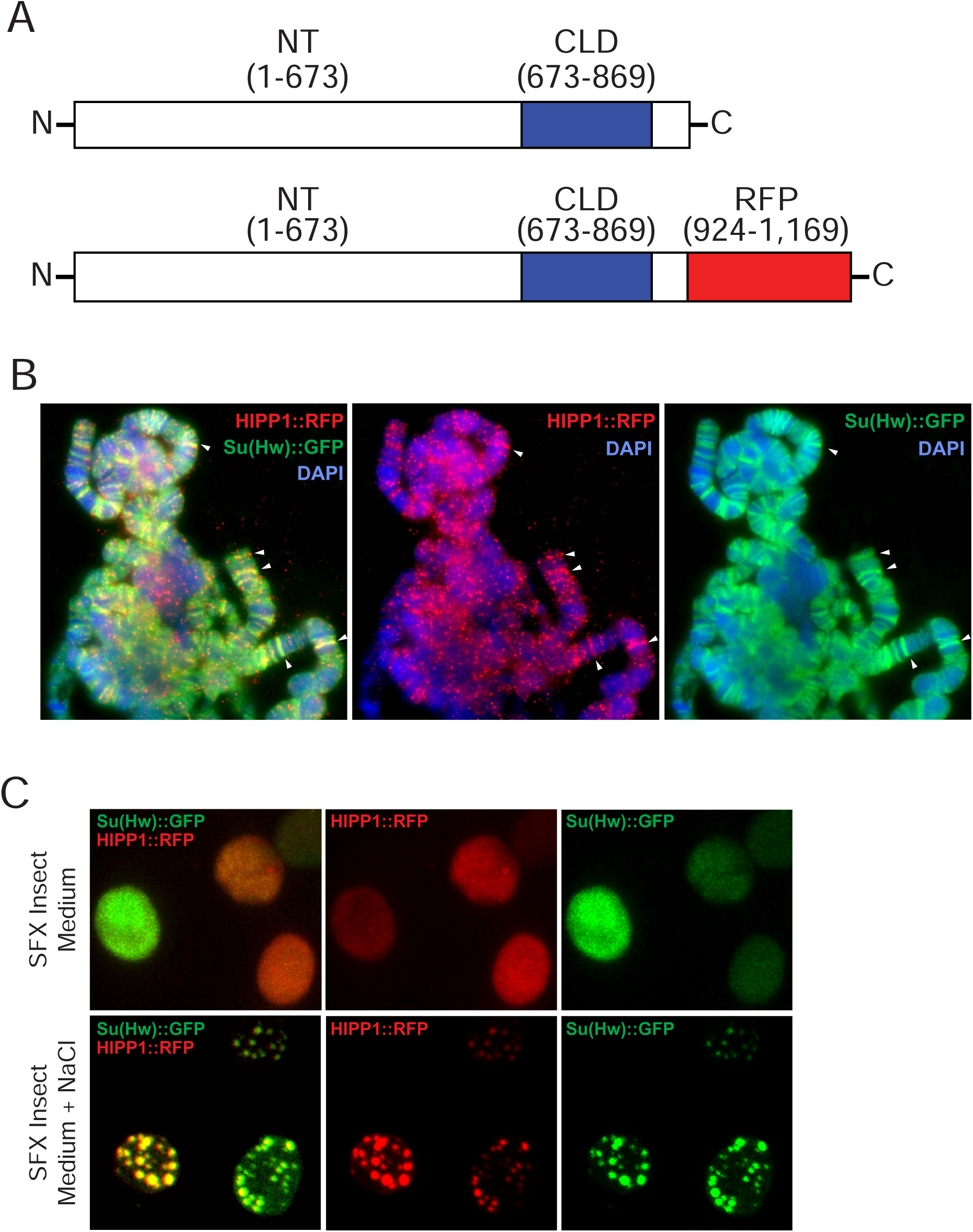
Su(Hw) and HIPP1 colocalize in polytene chromosomes and in insulator bodies during osmotic stress. (A) Diagrams of wild-type and transgenic HIPP1 are shown. The HIPP1 used for transgenic constructs in this work contains a C-terminal MCherry (mc) domain. (B) A polytene chromosome from larvae P{Su(Hw)::GFP, w^+^}, P{HIPP1::mc, w^+^}, vg-Gal4, labeled with anti-GFP and -RFP antibodies. Su(Hw)::GFP is shown in green, HIPP1::mc is shown in red, and DAPI is shown in blue. (C) S2 cells expressing transgenic Su(Hw)::GFP and HIPP1::mc transgenic constructs. Su(Hw)::GFP is shown in green and HIPP1::mc is shown in red. The top panel contains cells grown in normal media. The bottom panel shows cells treated for 20 minutes with media containing 250 mM added NaCl.

We also observed that the distribution of Su(Hw) and HIPP1 binding sites, relative to the band/interband structure of polytene chromosomes, was different among nuclei. In some nuclei Su(Hw) and HIPP1 localize exclusively to bands while in other nuclei they localize exclusively to interbands (Figure 3A and B). These observed changes in binding patterns suggest that Su(Hw) and HIPP1 binding is dynamic and is likely regulated in a cell cycle-specific manner. We also observe that some nuclei lack HIPP1::mc altogether (Figure 3C). This suggests that HIPP1 association with Su(Hw) is dynamically regulated and may contribute to a function of Su(Hw) only during specific stages of the cell cycle. The dynamic variability in HIPP1 and Su(Hw) binding patterns further suggests these proteins may be regulated by cell-cycle dependent processes. We hypothesize that the interaction between HIPP1 and Su(Hw) occurs transiently during the cell cycle, and that HIPP1 contributes to cell cycle-specific aspects of insulator activity such as regulating Su(Hw) function during DNA replication.

**Figure 3.**
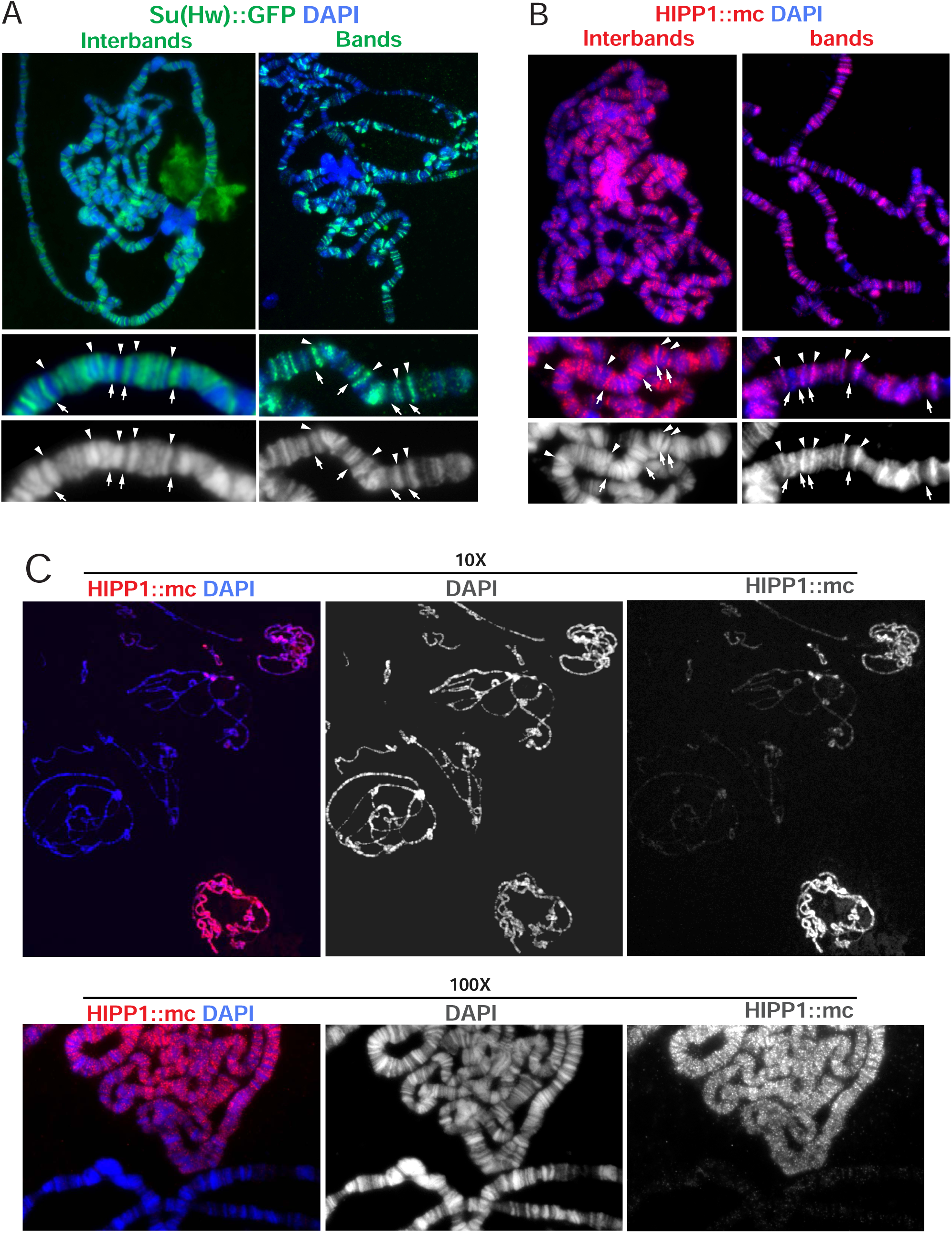
Binding patterns of Su(Hw) and HIPP1 to polytene chromosomes is variable in different nuclei suggesting may be dynamic and cell cycle-dependent. (A) Polytene chromosomes from P{Su(Hw)::GFP, w^+^}/vg-Gal4 larvae expressing Su(Hw)::GFP. Su(Hw), labeled with an anti-GFP antibody, localizes to either all interbands (left) or all bands (right) of the polytene chromosome. In zoom in images arrows point to interbands and arrowheads point to bands. (B) Polytene chromosomes from P{HIPP1::mc, w^+^}/vg-Gal4. HIPP1 larvae expressing HIPP1::mc. HIPP1 labeled with an anti-RFP antibody localizes to either all interbands (left) or all bands (right) of the polytene chromosome. Arrows and arrowheads as in A. (C) In addition to the exclusive band or interband binding pattern, many nuclei show no chromosome binding of HIPP1::mc. Polytene chromosomes from P{HIPP1::mc, w^+^}/vg-Gal4 immunostained with anti-RFP antibody at 10x (top) and 100x (bottom).

### Overexpression of Su(Hw) and HIPP1 disrupts cell proliferation

To explore the hypothesis that Su(Hw) may play a role in replication, we next determined whether driving overexpression of Su(Hw) and HIPP1 transgenic constructs impacts cell cycle progression. Driving the expression of Su(Hw)::GFP with a *vg*-Gal4 driver revealed significant cell proliferation defects in the adult wing margin, while driving the expression of HIPP1::mc with the same driver revealed no wing margin defects (Figure 4A and B, p=0.0003). Driving the expression of both Su(Hw)::GFP and HIPP1::mc in the same individuals resulted in wing margin defects that do not signifcantly differ from Su(Hw):: GFP expression alone (Figure 4A and B). To confirm that these defects in wing morphology were not due to apoptosis, we overexpressed p35 along with Su(Hw)::GFP (Figure 4A). p35 is a potent caspase inhibitor in *Drosophila* (Miller, 1997). Defects in the wing margin persisted, suggesting that the lack of cells in the wing margin are due to a lack of cell proliferation rather than apoptosis induced by elevated levels of Su(Hw) protein. Similar phenotypes are produced by mutations in the Notch pathway with also result in inhibition of cell proliferation in the wing margin (Baonza and Garcia-Bellido, 2000). These observations led us to conclude that Su(Hw) overexpression limits cell proliferation in the wing margin and that the lack of cell proliferation persists with the combined overexpression of Su(Hw)::GFP and HIPP1::mc. These data also shows that overexpression of HIPP1 does not rescue cell proliferation defects arising from Su(Hw) overexpression.

**Figure 4.**
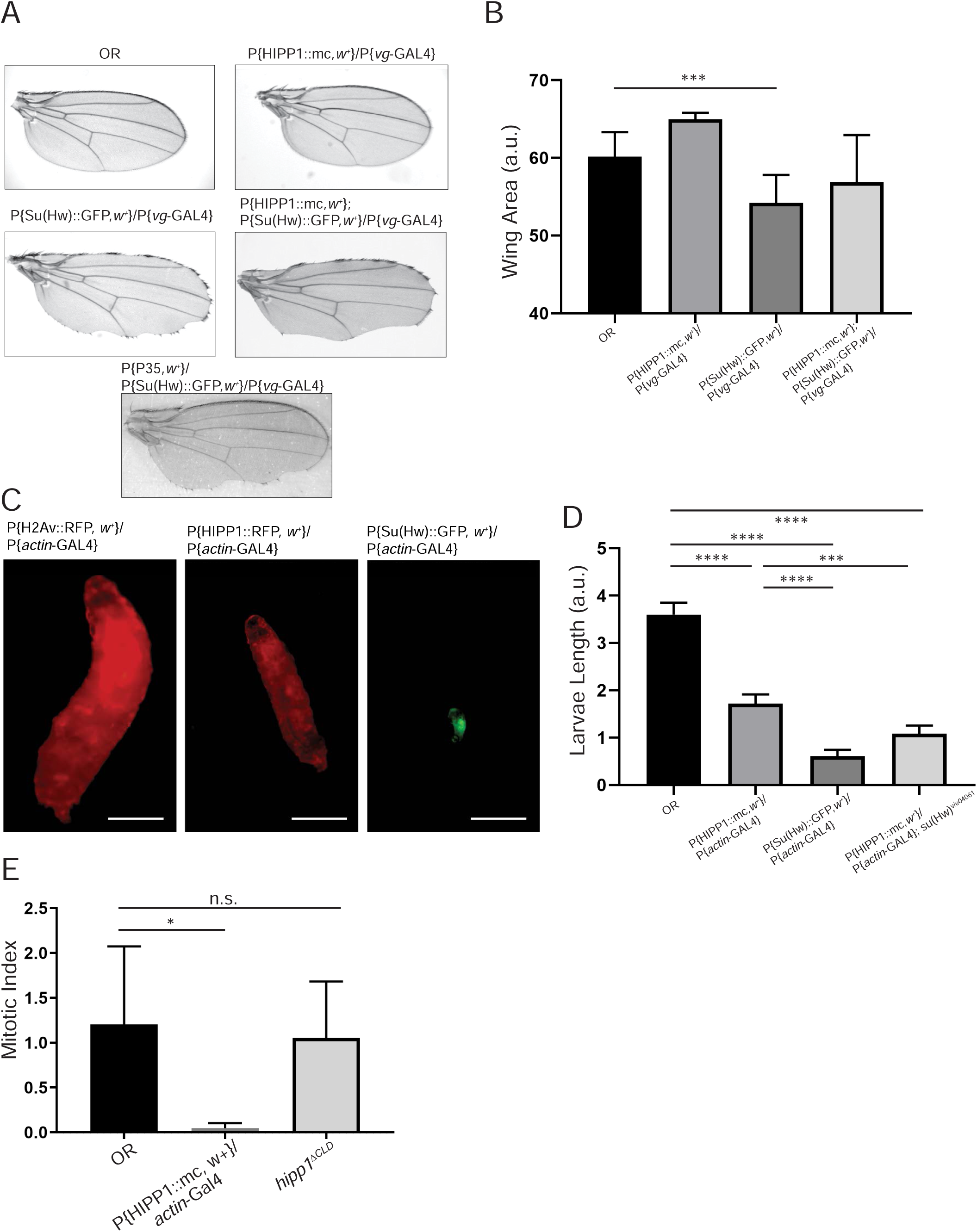
Su(Hw) and HIPP1 overexpression disrupts cell proliferation. (A) Images of wings from Oregon-R, P{Su(Hw)::GFP, w^+^}/vg-Gal4, P{HIPP1::mc, w^+^}/ vg-Gal4, or P{Su(Hw)::GFP, w^+^}; P{HIPP1::mc, w^+^}/vg-Gal4 flies are shown. Cuts in the wing margin suggesting cell proliferation defects can be seen in P{Su(Hw)::GFP, w^+^}/vg-Gal4 and P{Su(Hw)::GFP, w^+^}; P{HIPP1::mc, w^+^}/vg-Gal4 wings. (B) A bar graph quantifying the wing blade area from Oregon-R, P{Su(Hw)::GFP, w^+^}/vg-Gal4, P{HIPP1::mc, w^+^}/vg-Gal4, and P{Su(Hw)::GFP, w^+^}; P{HIPP1::mc, w^+^}/vg-Gal4 (***, p=0.0003). (C) Images showing the relative size of larvae, including control P{H2Av::mc, w^+^}/actin-gal4 over expressing H2Av::mc, P{Su(Hw)::GFP, w^+^}/ actin-Gal4, overexpressing Su(Hw)::GFP or P{HIPP1::mc, w^+^}/actin-Gal4 overexpressing HIPP1:mc. Scale bar: 2 mm (D) A bar graph of the measured lengths of Oregon-R, P{HIPP1::mc, w^+^}/actin-Gal4, and P{Su(Hw)::GFP, w^+^}/actin-Gal4 larvae (****, P<0.0001; ***, p=0.0006). (E) A graph showing the mitotic index, ratio of mitotic nuclei to total nuclei, for Oregon-R, P{HIPP1::mc, w^+^}/actin-Gal4, and *hipp1^ΔCLD^* larval brains. P{HIPP1::mc, w^+^}/actin-Gal4 brains show a significant reduction in the mitotic index, compared with Oregon-R (p=0.0410). P values were determined using an unpaired t-test.

Based on these observations, we tested how Su(Hw) and HIPP1 overexpression may affect *Drosophila* growth. Driving the expression of HIPP1::mc and Su(Hw)::GFP with a ubiquitous *actin-*Gal4 driver revealed a significant decrease in larval body size. We compared the sizes of HIPP1::mc and Su(Hw)::GFP overexpressing larvae to larva sizes from a line expressing H2Av::mc with the same *actin-*Gal4 driver. Both HIPP1::mc and Su(Hw)::GFP expression resulted in a decrease in larval body size, with Su(Hw)::GFP expression exhibiting a greater reduction in size (Figure 4 C and D), suggesting that Su(Hw) overexpression serves as a more potent inhibitor of cell proliferation compared to HIPP1. Both HIPP1::mc/*actin*-Gal4 and Su(Hw)::GFP/*actin*-Gal4 larvae die without reaching sizes larger than shown in Figure 4, and never enter pupation stage.

We additonally measured growth in larve expressing HIPP1::mc with an *actin*-Gal4 driver in a *su(Hw)^v/e041061^* mutant background. These larvae exhibit a reduced size, significantly smaller than HIPP1 overexpression alone. The inability of mutations in *su(Hw)* to rescue the effects of HIPP1 overexpression on larval growth suggests roles for HIPP1 in cell proliferation that extend beyond interactions with the Su(Hw) insulator complex alone. HIPP1 interacts with other protein complexes including CTCF and HP1 (Alekseyenko *et al*., 2014). Therefore, HIPP1 overexpression may contribute to cell proliferation defects through interactions with CTCF and HP1, as well as through interactions with Su(Hw). It is also of note that the combination of HIPP1 overexpression and mutations in *su(Hw)* do not rescue the small larvae phenotype and, instead, causes a significant reduction in size compared to HIPP1 overexpression alone. Mutations in *su(Hw)* have been linked to developmental defects and replication stress (Hsu *et al*., 2019; Klug *et al*., 1968; Klug *et al*., 1970). These results suggest that HIPP1 overexpression combined with mutations in *su(Hw)* promote more severe cell proliferation defects compared to HIPP1 overexpression alone. Next, we measured the mitotic index of dividing neuroblasts in larval brains expressing HIPP1::mc to further assess whether defects observed in wing development and larval growth are the result of cell cycle disruption. We measured the mitotic index in HIPP1::mc/*actin-*Gal4 larval brains and found they have a significantly lower mitotic index compared to the Oregon-R control (Figure 4E, p=0.0410). This result suggests HIPP1-overexpressing cells complete the cell cycle less frequently that Oregon-R cells, causing insufficient cell proliferation and providing an explanation for the reduced size previously noted in HIPP1 overexpressing larvae.

### HIPP1 overexpression delays the transition of DNA replication between early and late replicating regions of the genome

Our observations of delays in the cell cycle when Su(Hw) and HIPP1 proteins are overexpressed in the nucleus suggests that these proteins serve as barriers to normal cell cycle progression. The association of Su(Hw) with the nuclear lamina and regions flanking intercalary heterochromatin domains suggests Su(Hw) plays a role in maintaining functional boundaries between LADs and actively transcribed TADs in the nuclear interior (Khoroshko *et al*., 2016). One possibility is that in addition to functioning as a boundary, the Su(Hw) insulator complex also mediates the detachment of the insulator from LADs and from euchromatin/heterochromatin transition sites, thereby facilitating access into these domains by cellular machinery during genome replication. Since heterochromatin domains flanked by Su(Hw) binding sites should be late replicating domains, we asked whether changes in Su(Hw) or HIPP1 expression levels alter the rate of DNA replication within each domain.

Due to the cell proliferation defects observed in HIPP1-overexpressing conditions, we hypothesized that HIPP1 may affect the progression of replication forks when present in its overexpressed form. To test this hypothesis, we quantified the amount of single strand DNA (ssDNA) in Oregon-R, *hipp1^ΔCLD^*, and HIPP1::mc/*actin*-Gal4 larval brains using ssDNA as a marker for active replication forks (Zellweger *et al*., 2015) (Figure 5A). BrdU is a nucleotide analog used to monitor nucleodtide incorporation during DNA replication. The detection of BrdU incorporation into DNA relies on the binding of a BrdU-specific antibody. The anti-BrdU antibody, however, can only detect incorporated BrdU within the DNA if the DNA is single stranded, thus DNA must be denatured prior to antibody incubation. Detecting BrdU incorporation in non-denaturing conditions provides a way to measure the amount of ssDNA present at stalled or active replication forks (Despras *et al*., 2010). We incubated Oregon-R, *hipp1^ΔCLD^*, and HIPP1::mc/*actin*-Gal4 larval brains with BrdU for one hour, followed by fixation and immunostaining. We found a significant increase in BrdU accumulation in HIPP1 overexpression larval brains compared to the Oregon-R control (Figure 5B). This suggests that overexpression of HIPP1 leads to an accumulation of active and stalled replication forks. Taken together with our observation that HIPP1 overexpression causes a decrease in the mitotic index, the observed accumulation of ssDNA suggests that HIPP1 overexpression disrupts DNA replication, possibly leading to the activation of checkpoints and stalling of the cell cycle. We hypothesize that the interaction between HIPP1 and Su(Hw) alters insulator activity in a cell cycle-specific manner and that HIPP1 overexpression prolongs this change in insulator function, thereby misregulating insulator properties and possibly aspects of the replication timing.

**Figure 5.**
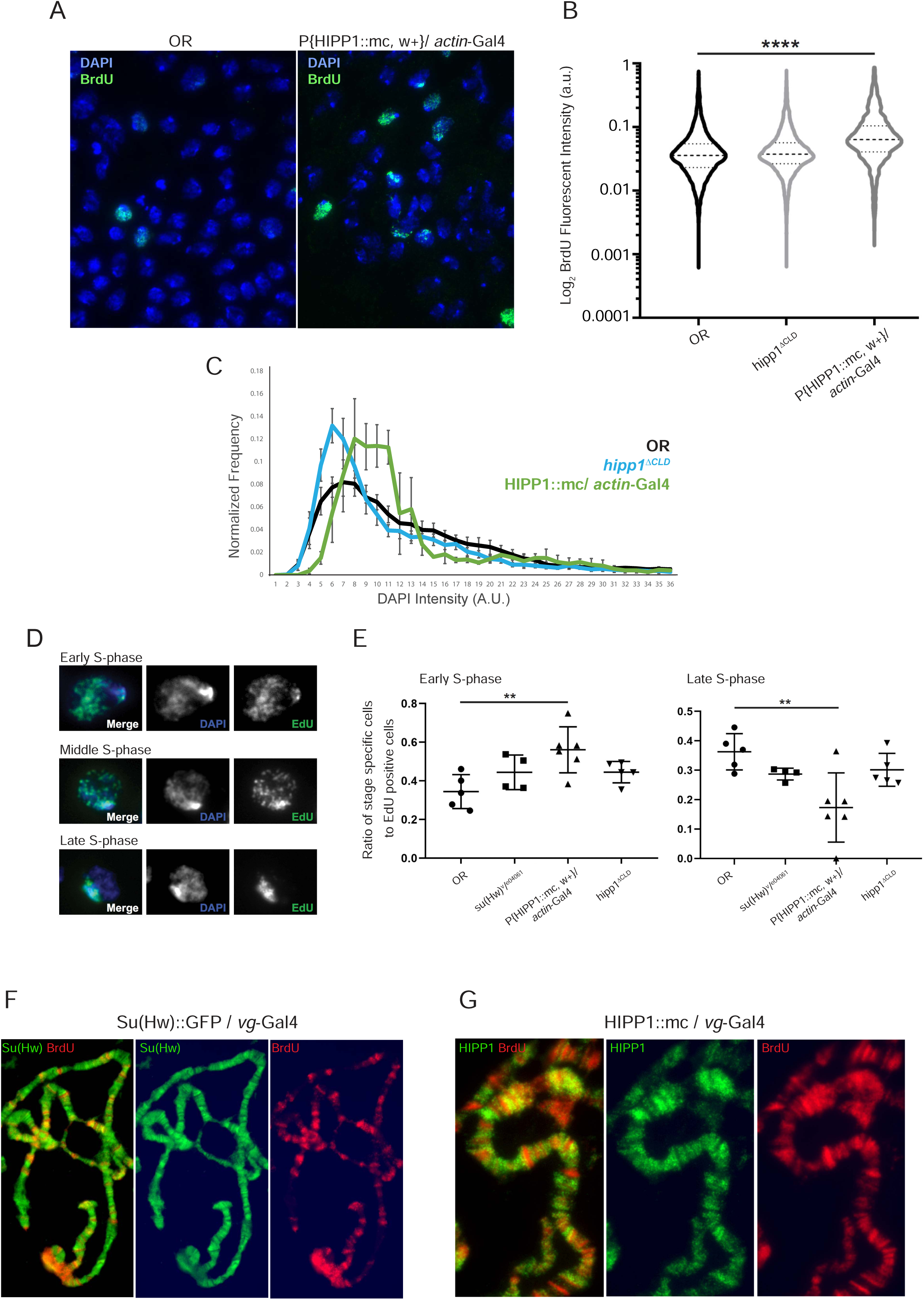
HIPP1 overexpression alters replication timing in larval brain cells. (A) Representative images of brain cells from Oregon-R and P{HIPP1::mc, w^+^}/actin-Gal4 larvae labeled with DAPI (blue) and incorporated BrdU (green). (B) Quantification of BrdU fluorescent intensity per nuclei in Oregon-R, *hipp1^ΔCLD^*, and P{HIPP1::mc, w^+^}/actin-Gal4 larval brain cells. (P<0.0001). (C) Frequency of nuclei with different levels of DAPI intensity from Oregon-R, *hipp1^ΔCLD^*, and P{HIPP1::mc, w^+^}/actin-Gal4 larval brain cells are charted. P{HIPP1::mc, w^+^}/ actin-Gal4 larval brain cells exhibit a shift towards a higher level of DAPI intensity. (D) Representative images of larval brain cells labeled with DAPI (blue) and EdU (green) in Early, Middle, and Late S-phase. (E) P{HIPP1::mc, w^+^}/actin-Gal4 larval brains have a significantly larger ratio of cells in the early replication phase, compared to Oregon-R (p=0.0073). *su(Hw)^e04061/v^* mutants, and *hipp1^ΔCLD^* mutants were also considered. P{HIPP1::mc, w^+^}/actin-Gal4 larval brains have a significantly lower ratio of cells in the late replication phase, compared to Oregon-R (p=0.0095). P values were determined using an unpaired t-test. (F) Polytene chromosomes from P{Su(Hw)::GFP, w^+^}/vg-Gal4 larvae showing Su(Hw)::GPP in green, labeled using anti-GFP antibody, and BrdU in red, labeled with anti-BRuU. BrdU incorporation is enriched at sites of low Su(Hw)::GFP staining. (G) Polytene chromosomes from P{HIPP1::mc, w^+^}/vg-Gal4 larvae showing HIPP1::mc in green, labeled with anti-RFP antibody, and BrdU in red. BrdU incorporation is enriched at sites of low HIPP1::mc staining.

Next, we quantified the level of DAPI intensity in the larval brain samples from our BrdU experiments to observe differences in cell cycle stages. DAPI intensity is a common method to determine phases of the cell cycle (Stohr *et al*., 1977). Cells in G1 contain only one copy of each chromosome and cells in S and G2 contain greater than one copy of each chromosome. By quantifying the frequency of cells that contain different ranges of DAPI intensity, we observed that HIPP1::mc/*actin*-Gal4 cells were biased towards a greater DAPI content, compared to Oregon-R or *hipp1^ΔCLD^* cells (Figure 5C). This data suggests that a large fraction of HIPP1::mc/*actin*-Gal4 cells are arrested during S or G2 phases, possibly a consequence of replication fork stalling. We also observe a shift towards less DNA content in *hipp1^ΔCLD^* cells, suggesting replication is suppressed in *hipp1^ΔCLD^* mutants, potentially through a Su(Hw)-dependent mechanism (Figure 5C).

To further investigate the influence of Su(Hw) and HIPP1 on DNA replication, we used 5-ethynyl-2-deoxyuridine (EdU) as a marker for DNA replication to monitor the progression of DNA replication in HIPP1-overexpressing organisms. EdU is a thymidine analog that incoporates during DNA replication and can be detected by activation of an EdU-specific label. Incubation of tissues for a fixed amount of time allows us to observe the number of cells undergoing S-phase as well as the genomic location of active DNA replication. We incubated larval brains in EdU for ten minutes including brains dissected from Oregon-R, *su(Hw)^v/e04061^*, *hipp1^ΔCLD^*, and HIPP1::mc/*actin-*Gal4 larvae (brains from Su(Hw)::GFP/*actin-*Gal4 larvae do not develop to a large enough size to allow dissection). We then detected EdU incorporation using an EdU-specific label and fluorescence microscopy (Figure 5D). DNA replication occurs in distinct phases dependent upon chromatin state (Armstrong *et al*., 2018; Lubelsky *et al*., 2014). Euchromatin replicates during early S-phase, a combination of euchromatin and heterochromatin replicate during middle phase, and constitutive heterochromatin replicates during late S-phase. Based on EdU labeling of larval brains, we quantified the number of cells in each phase for each genotype. We found that there are singificantly more HIPP1::mc/*actin-*Gal4 cells in the early S-phase, compared to Oregon R (Figure 5E, Early S-phase, p=0.0073). We also found that significantly fewer HIPP1::mc/*actin-*Gal4 cells were in the late replication compared to Oregon-R (Figure 5E, Late S-phase, p=0.0095). This result suggests that HIPP1 overexpression stalls progression of DNA replication in the early replication phase and delays the entry of replication machinery into middle and late replicating regions. Additionally, we noticed no significant change in *su(Hw)^v/e04061^* or hipp1*^ΔCLD^* mutants. This suggests that the delay in the replication timing program is dependent upon HIPP1 overexpression.

We next aimed to observe nucleotide incorporation at a higher resolution to better understand the slow progression of S-phase in HIPP1::mc/*actin-*Gal4 cells. To do so, we visualized the incorporation of 5-Bromo-2’-deoxyuridine (BrdU) relative to Su(Hw) and HIPP1 in the salivary glands of HIPP1::mc/*vg-*Gal4 and Su(Hw):: GFP/*vg-*Gal4 organisms. We found that in some genomes both Su(Hw)::GFP and HIPP1::mc signals occurred opposed to BrdU signal. This pattern of BrdU incorporation suggests that HIPP1 and Su(Hw) may be barriers to DNA replication at certain stages during genome replication, possibly slowing down the S-phase and cell proliferation when overexpressed (Figure 5F).

To better understand the effect of Su(Hw) and HIPP1 on the progression of S-phase, we next observed the time that it takes for Oregon-R, *hipp1^ΔCLD^*, HIPP1::mc/*vg*-Gal4, and Su(Hw):: GFP/*vg*-Gal4 to complete each stage of the S-phase using polytene chromosomes from third instar larvae salivary glands. Polytenes undergo endocycling in which they only participate in G and S-phases (Smith and Orr-Weaver, 1991). Chromatin is distributed into condensed heterochromatin-like bands and euchromatin interbands along the arms of polytene chromosomes, with pericentric heterochromatin located at the chromocenter. We timed stages of S-phase in polytenes by incubating salivary glands in cell culture media for periods of time, followed by fixation and antibody staining. Specifically, we incubated salivary glands with EdU for 10 minutes at the beginning of each experiment. We then washed the EdU from salivary glands and allowed them to incubate in media for 2, 4, 6 or 8 hours. Following incubation, tissues were immediately fixed and stained for PCNA. By comparing areas of EdU incorporation with areas of PCNA immunostaining, we were able to estimate a progression time between different S-phase stages in each of the lines. For instance, if a polytene displayed EdU in early/late phase pattern and PCNA in an end of S-phase pattern after 6 hours of incubation, we concluded that it took the chromosome 6 hours to progress from the early/late phase of replication to the end of S-phase replication stage (Figure 6A). We assigned EdU and PCNA staining patterns to phases of DNA replication according to a previous study (Kolesnikova *et al*., 2013). This study labeled polytene chromosomes with PCNA and characterized early S-phase (I) as continuous PCNA signal, early to late phase (II) as PCNA signal in polytene arm bands, late S-phase (III) as PCNA in the chromocenter and intercalary heterochromatin, and the end of S-phase (IV) as very weak PCNA signal in the chromocenter and intercalary heterochromatin. Stage V is assigned to chromosomes in G phase and show no PCNA signal. Under these conditions, we found that HIPP1::mc/*vg*-Gal4 chromosomes require significantly more time to progress between replication phases when compared with Oregon-R cells. This result further suggests HIPP1 plays a role in DNA replication and that misregulation of HIPP1 by overexpression causes replication to progress more slowly (Figure 6A and B).

**Figure 6.**
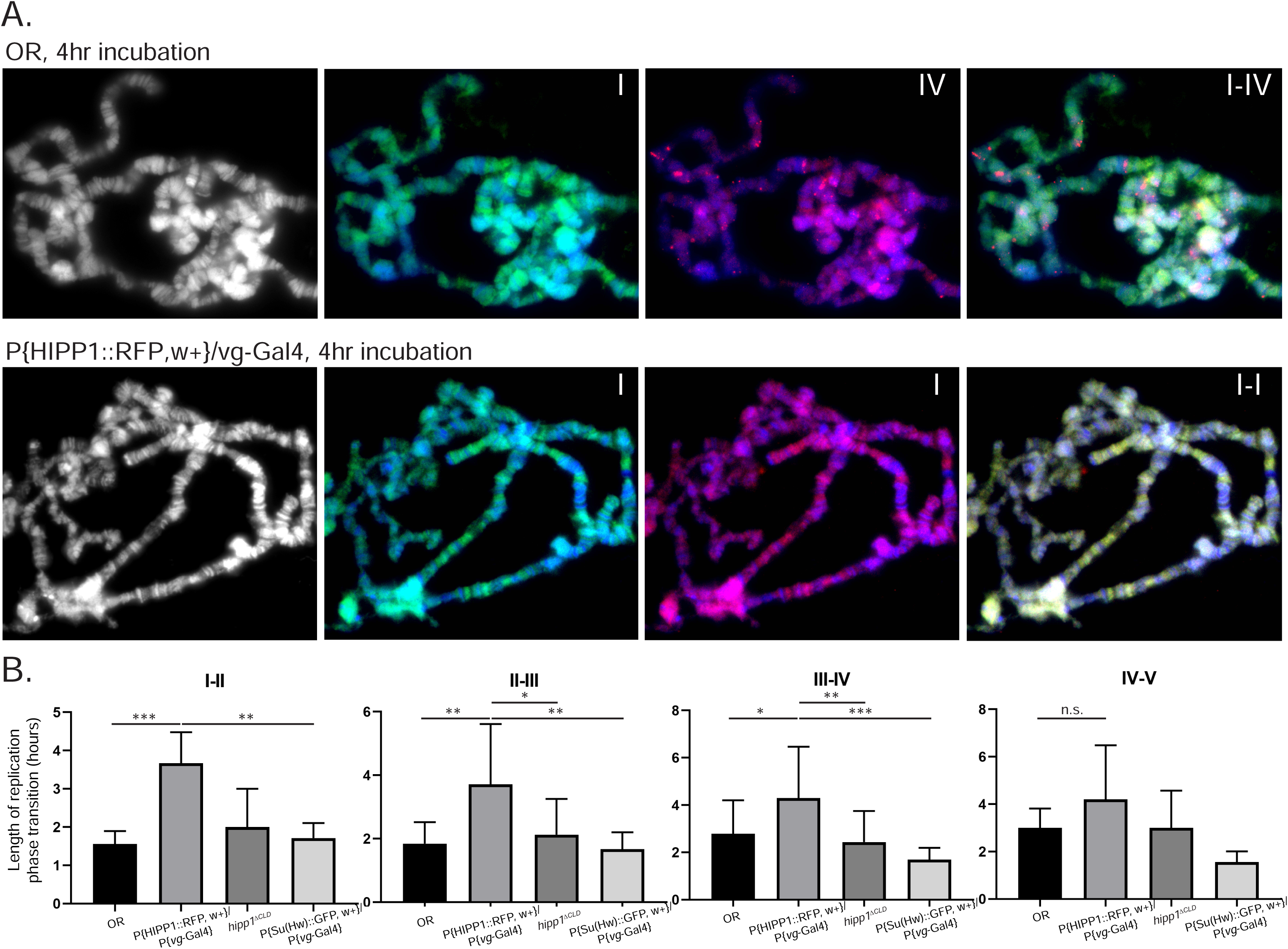
HIPP1 overexpression delays the progress of replication in polytene chromosomes. (A) Images of polytene chromosomes from Oregon-R (top) and P{HIPP1::mc, w^+^}/vg-Gal4 (bottom) salivary glands. Chromosomes were initially labeled with EdU, followed by fixation and staining with an anti-PCNA antibody after certain time points, specifically 2, 4, 6, and 8 hours after the EdU incubation. In the example, Oregon-R chromosomes progress from early S-phase (I) to the end of S-phase (IV) in 4 hours. P{HIPP1::mc, w^+^}/vg-Gal4 chromosomes remain in the early S-phase (I) after the 4-hour incubation. (B) Quantification of how many chromosomes from each time point were able to complete each phase transition (I-II, II-III, III-IV, IV-V). P values using an unpaired t-test are as follows: I-II, ***, p=0.0007, **, p=0.0011; II-III, **, p=0.003, *, p=0.0230, **, p=0.0014; III-IV, *, p=0.0199, **, p=0.006, ***, p=0.0001.

### HIPP1 overexpression disrupts Su(Hw) insulator function

Together, our results suggest that the effects of HIPP1 on cell cycle progression could be both dependent and independent from Su(Hw) insulators. This conclusion is not unexpected since, in addition to Su(Hw), HIPP1 interacts with other insulator proteins such as dCTCF, as well as with HP1. In each one of these interactions HIPP1 may have a putative role on the progression of the cell cycle. To uncover the role of HIPP1 in insulator function, we analyzed Su(Hw)-dependent phenotypes in flies overexpressing HIPP1. The *yellow^2^* and *cut^6^* mutations (*y^2^*and *ct^6^*) are caused by an insertion of the *Gypsy* retrotransposon between tissue specific enhancers and the promoter of these genes, allowing the Su(Hw) insulator protein to bind and disrupt normal enhancer promoter interactions and gene transcription activation (Jack *et al*., 1991; Parkhurst and Corces, 1986). These mutations result in flies with yellow body and yellow wings (*y^2^*), and cuts in the wing margin (*ct^6^*). Interestingly, we observed that driving HIPP1 overexpression with a *vestigial*-Gal4 promoter in flies with *y^2^ct^6^*background results in the restoration of wild-type black wing blades and rounded wing margins (Figure 7A). Immunostaining of polytene chromosomes from larvae with the same genotype reveal that Su(Hw) remains bound to the *yellow* locus in HIPP1-overexpressing polytene chromosomes, despite the suppression of insulator function (Figure 7B). This result suggests overexpression of HIPP1 suppresses the enhancer blocking activity of the Su(Hw) insulator without disrupting the binding of Su(Hw) to DNA. We also observe that *y^2^ct^6^*; *hipp1^ΔCLD^* mutants display no rescue of the *y^2^* and *ct^6^* phenotypes, suggesting that the activity of the CLD is not necessary for insulator function (Figure 7A).

**Figure 7.**
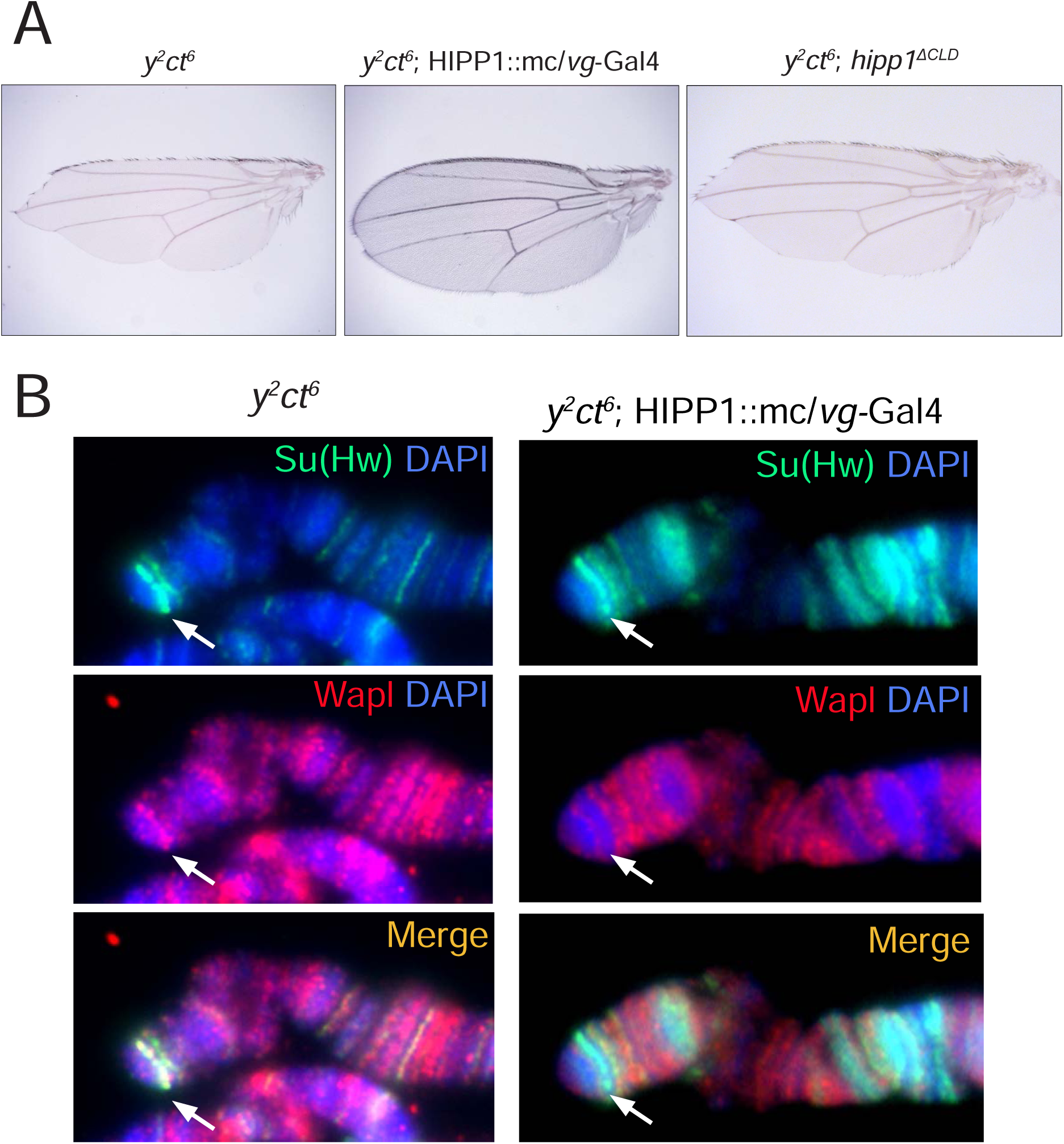
HIPP1 overexpression rescues *y^2^* and *c^6^*phenotypes. (A) *y^2^ct^6^*, *y^2^ct^6^*; P{HIPP1::mc, w^+^}/vg-Gal4, and *y^2^ct^6^*; *hipp1^ΔCLD^* wings are shown. Only wings from *y^2^ct^6^*; P{HIPP1::mc, w^+^}/ vg-Gal4 flies look phenotypically normal, showing almost perfectly round margins and black blades, indicating that overexpression of HIPP1::mc suppresses gypsy enhancer-blocking activity. (B) Polytene chromosomes from *y^2^ct^6^* and a *y^2^ct^6^*; P{HIPP1::mc, w^+^}/vg-Gal4 larvae, showing the tip of the X chromosome and the Su(Hw) band on the *y^2^ gypsy* insertion. Su(Hw) is shown in green, WAPL is shown in red, and DAPI is shown in blue. Su(Hw) and WAPL colocalize at the *y^2^* site (arrow). Su(Hw) and WAPL do not colocalize at the *y^2^* site in the *y^2^ct^6^*; P{HIPP1::mc, w^+^}/ vg-Gal4 chromosome (arrow).

*Drosophila* insulators lack a comprehensive model that combines the canonical roles of insulator proteins such as enhancer blocking and boundary formation with genome-wide organizational properties such as loop formation. Recent computer modeling and work in mammalian systems have provided a model of loop extrusion by cohesins to explain the multifaceted role of mammalian CTCF in shaping the genome while contributing to gene transcription (Fudenberg *et al*., 2016). The presence of loop extrusion-like domains in *Drosophila* has been disputed (Rowley *et al*., 2017), but recent work analyzing Hi-C maps of the *Drosophila* genome at a resolution of ∼200 bp reveals the presence of TADs defined by insulator binding sites, suggesting the same organizational principles are conserved between insects and mammals (Wang *et al*., 2018). These advances in elucidating the involvement of the cohesin complex in driving loop formation and the confirmation of TAD organization in *Drosophila* motivated us to further explore whether the cohesin complex plays a role in the insulator function of the gypsy insulator.

Using fluorescence microscopy in polytene chromosomes, we observed that WAPL, a component of the cohesin complex, colocalizes with Su(Hw) and other insulators at a number of sites including *y^2^* sites (Cunningham *et al*., 2012) (Figure 7B). However, in HIPP1::mc/*vg*-Gal4 overexpression conditions, we observe that WAPL no longer colocalizes with Su(Hw). Although the formation of loops mediated by cohesin and insulators have not directly been observed in *Drosophila*, similarities between organizational properties in human and *Drosophila* genomes point towards a conserved mechanism. We hypothesize that cohesin contributes to the insulator function of Su(Hw). Under this assumption, when HIPP1 is overexpressed and WAPL leaves the insulator site, Su(Hw) is no longer able to act as an insulator. Altogether these data suggest HIPP1 modulates replication timing by regulating insulator activity in a cell cycle- or genome replication-dependent manner.

## Discussion

Insulator binding sites are abundant in the genome and play critical roles in genome structure and function. Null mutations in mammalian CTCF and in most insulator proteins in *Drosophila*, including dCTCF, often result in lethality. Therefore, it is surprising that mutations in *su(Hw)* have no discernible effect during development or in the adult organism other than female sterility (Hsu *et al*., 2015; Klug *et al*., 1968; Klug *et al*., 1970; Soshnev *et al*., 2013). Interestingly, a null mutation of HIPP1 is also viable with no discernible phenotype in adults (Glenn and Geyer, 2018). This raises the question of whether these proteins truly play a role in important 3D genome organization and function, in addition to contributing to tissue specific gene transcription regulation. Results presented here and elsewhere show that mutant and overexpressing genotypes of Su(Hw) and HIPP1 generate phenotypes related with defects in DNA repair and genome replication (Hsu *et al*., 2019).

Coinciding with the findings of Glenn and Geyer (2018), we found that the CLD specific mutant produced in this report (*hipp1^ΔCLD^*) also has no obvious effect on development or insulator function. However, we do find that *hipp1^ΔCLD^* has an impaired ability to repair double-strand breaks produced following irradiation treatment. Glenn and Geyer (2018), previously identified HIPP1 as the *Drosophila* homolog of the human CDYL protein. However they found that *hipp1* mutants did not have the same consequences on viability and male fertility as mutants for human *cdyl* (Glenn and Geyer, 2018). Here, we investigated whether the CLD of HIPP1 and CDYL share a role in DNA repair. The CLD of human CDYL is an important component of the homology directed repair pathway (Abu-Zhayia *et al*., 2018). We find that *hipp1^ΔCLD^* mutants display an impaired response to DNA damage. Additional work will be required to establish to what extent the role of HIPP1 in DNA repair in *Drosophila* is conserved with that of CDYL in humans. Although a connection between the Su(Hw) insulator complex and DNA damage repair has not yet been established, it has been reported that the CTCF boundary function in humans is linked to double-strand break formation by topoisomerase activity (Canela *et al*., 2017). Additionally, it has been shown that topoisomerase II modulates Su(Hw) insualtor function in *Drosophila* (Ramos *et al*., 2011). This evidence suggests the possibility that insulator binding sites in *Drosophila* may also be sites that accumulate torsional stress, which may lead to replication forks stalling during DNA replication and rely on HIPP1 for efficient torsion relief and/or repair.

Supporting these observations, we also find that WAPL, a component of the cohesin complex, colocalizes with the Su(Hw) insulator complex at the *y^2^* locus. The cohesin complex has been identified as a critical component of CTCF-mediated loops in mammalian systems and through the loop extrusion mechanism is thought to be responsible for accumulation of torsional stress a CTCF sites in mammals (Canela *et al*., 2017). Our finding that WAPL colocalizes with the *gypsy* insulator suggests insects have a conserved mechanism of insulator mediated DNA looping (Fudenberg *et al*., 2016). Our results show that upon HIPP1 overexpression, *y^2^* and *ct^6^*phenotypes are reversed to normal while simultaneously WAPL no longer colocalizes with the corresponding Su(Hw) insulator sites. The correlation between WAPL presence and the enhancer-blocking activity of the Su(Hw) insulator suggests cohesin plays a role in stabilizing the interactions required for enhancer-blocking in the Su(Hw) insulator complex. These observations indicate that the role of HIPP1 binding to Su(Hw) may be transient, i.e. its role is to temporarily destabilize cohesin’s association with the insulator complex and transiently suppress the compartment boundary activity of the insulator.

Our data shows that an increase in HIPP1 expression both inhibits phenotypes mediated by the Su(Hw) insulator and delays cell cycle progression. These findings, taken together with our observations that overexpression of Su(Hw) and HIPP1 delays cell cycle progression suggest that HIPP1 antagonizes insulator function as a part of a mechanism that regulates progression of replication forks through different genome compartments. Earlier studies establishing the association of Su(Hw) with sites flanking intercalary heterochromatin and the initial observation that Su(Hw) colocalizes with multiple replication origins directed our attention into the question of whether the Su(Hw)-HIPP1 interaction may play a role in DNA replication (Khoroshko *et al*., 2016; Vorobyeva *et al*., 2013). We observe that Su(Hw) overexpression serves as a potent inhibitor of cell proliferation, causing a greater decrease in cell proliferation than HIPP1 overexpression. Additionally, we find that HIPP1 overexpression has a negative effect on DNA replication, resulting in an accumulation of early replicating cells and fewer late replicating cells. Our analysis of DNA replication in polytene chromosomes reveals that HIPP1 overexpression specifically delays the transition between stages of replication. These findings suggest that the transition between replicating domains is sensitive to the levels of HIPP1 expression. Since Su(Hw) binding sites are enriched in regions flanking intercalary heterochromatin, or late replicating domains, we hypothesize that HIPP1 modulates Su(Hw) insulator activity during S-phase perhaps by timing the replication entry into chromatin domains guarded by Su(Hw) binding sites. Higher amounts of HIPP1 may cause an imbalance in this process and result in misregulation of the transition between euchromatin and heterochromatin by replication machinery. The replication defects caused by HIPP1 overexpression, however, are quite dramatic and most likely cannot be accounted for by HIPP1-Su(Hw) interactions alone. HIPP1 has a variety of binding partners other than Su(Hw), including other insulator proteins such as dCTCF and HP1 (Alekseyenko *et al*., 2014). Future experiments are needed to address the mechanistic details of the relationship between HIPP1 and HP1 and other insulators.

The evidence presented here suggests HIPP1 interacts with Su(Hw) to regulate yet unknown aspects of replication timing in the *Drosophila* genome, but would appear to contradict recent findings demonstrating that deletion of insulator sites in mouse embryonic stem cells does not significantly affect compartmentalization of active and inactive regions of the genome or replication timing (Sima *et al*., 2019). Moreover, this finding would appear to be in line with the observations that Su(Hw) and HIPP1 null mutations are viable, or that dCTCF mutants, although ultimately lethal, allow for full embryo development in *Drosophila* (Gambetta and Furlong, 2018). On the other hand, detailed studies analyzing genome structure during development have identified a role for DNA replication in the establishment and maintenance of TADs by demonstrating that inhibition of DNA replication, rather than inhibition of transcription, prevents TAD formation during early mouse embryogenesis, further suggesting a link between replication programs and genome structure (Ke *et al*., 2017). In this context, a role of genome structure in normal genome replication is also reinforced by our previous finding that mutations in *su(Hw)* lead to replication stress in nurse cells and dividing neuroblasts (Hsu *et al*., 2019). The observation that Su(Hw)-deficient cells present replication defects challenges the accepted notion that *su(Hw)* null mutations allow for complete normal development. It is possible that the replication defects originating from the lack of *su(Hw)* alone are insufficient to prevent development. Perhaps it is the combination of multiple architectural proteins in *Drosophila*, or a combination of functionally different CTCF sites in mammals, which collectively shape the structure of the genome, that is needed for normal replication progression through genome compartments and normal replication timing.

In summary, here we have presented evidence that a Su(Hw)-interacting protein has the ability to regulate insulator activity and alters the rate of cell proliferation and the replication timing program when ectopically expressed. We propose that HIPP1 is a regulator of insulator activity. When the Su(Hw) insulator sites are knocked out genome-wide, flies are viable but actively replicating cells undergo replication stress, which suggest insulator function is required for normal replication timing (Hsu et al., 2019). When HIPP1 is overexpressed, we observe misregulation of genome replication and replication timing, possibly resulting from the ectopic inactivation of insulator function. These findings support the notion that genome replication is supported by a mechanism in which insulator function must be regulated to allow normal replication timing and cell-cycle progression.

## Materials and Methods

### Drosophila stocks

All fly stocks and crosses were maintained on standard cornmeal-agar media and yeast in a 25°C incubator. The fly stocks used in this work included: microinjection to generate transgenic lines *yw*; P{HIPP1::mc, *w^+^*}, *yw*; P{SuHw::EGFP, *w^+^*} (Schoborg *et al*., 2013), and *yw*; P{H2Av::mc, w^+^} were performed by GenetiVision; the lines obtained from the *Drosophila* Bloomington Stock Center at Indiana University: *su(Hw)^e041061^*/*TM6B;* w^+^; P{GAL4-vg.M}2; *TM2*/*TM6B*; the lines from V. Corces (Emory University): *su(Hw)^v^/TM6B, Tb1*, *mod(mdg4)^u1^*. The mutant lines *hipp1^ΔCLD31.2^* and *hipp1^ΔCLD14.3^* were generated by our lab using CRISPR Cas-9; microinjection of guide RNAs was performed by Gentivision.

### Antibodies

Rabbit anti-Su(Hw) polyclonal IgG antibody was generated in our laboratory (Wallace *et al*., 2010). Rabbit anti-WAPL polyclonal IgG was generated in the laboratory of Dr. Judith Kassis and used according to prior reports (Cunningham *et al*., 2012). The following commercially available primary antibodies were used: Mouse anti-GFP IgG (Developmental Studies Hybridoma Bank #12A6), rabbit anti-RFP IgG (A00682, GenScript), mouse anti-PCNA IgG (Abcam ab29), and mouse anti-BrdU IgG (Developmental Studies Hybridoma Bank G3G4). Antibodies were used at a concentration of 5 µg/ml. The following secondary antibodies were used: Donkey FITC-conjugated anti-mouse IgG (Jackson ImmunoResearch Laboratories, Inc.), Donkey Alexa Fluor 488-conjugated anti-rabbit IgG (A-21206, Life Technologies), and Donkey Alexa Fluor 488-conjugated anti-mouse IgG (A-21202, Thermo Fisher).

### Expression vector construction

Expression vectors for S2 cells and P-elements were created as previously described (Schoborg *et al*., 2013). The S2 cell dual-expression constructs contain Su(Hw)-EGFP and HIPP1-mCherry sequences, including introns, under the control of the copper-responsive metallothionein promoter in the pMK33-CTAP tag vector backbone. Fly expression constructs were created as previously described (Schoborg *et al*., 2013). Su(Hw)-EGFP and HIPP1-mCherry were amplified from pMK33 and inserted into the pUAST-Y vector backbone.

### Polytene chromosomes immunostaining and quantification

Salivary glands from early third instar larvae were dissected in insect media (HyClone SFX; Thermo Fisher Scientific), and fixed immediately with 4% PFA; 50% acetic acid on a coverslip. Salivary glands were squashed on a microscope slide until the polytene chromosomes were spread out. Slides were dipped in liquid nitrogen to remove coverslips. Polytene chromosomes were blocked for 10 minutes at room temperature (RT) in blocking solution (PBS, 0.1% NP40, 3% nonfat milk). Primary antibodies were diluted in blocking solution to a concentration of 5 µg/ml and incubated overnight at 4 °C in a humidified chamber. Primary antibodies were removed by incubating in washing buffer (PBS, 0.1% NP40) for 10 minutes at room temperature. Secondary antibodies were then diluted in blocking solution (1:200) and incubated for 1 hour at room temperature and washed as described before. DAPI (4’, 6-diamidino-2-phenylindole 0.5 µg/ml) was used to counter stain the DNA for 30 seconds before rinsing with PBS. Slides were mounted with Vectashield mounting medium (Vector Laboratories) and sealed with nail polish.

### Stress treatment and immunostaining

S2 cells 3–5 days after subculture were allowed to adhere to a poly-l-lysine coverslip for 30 min in a covered 35 mm cell culture dish. To induce osmostress, media was removed and quickly replaced with fresh SFX media supplemented with the indicated concentration of NaCl (from a 5M stock). Controls were kept in conditioned media. Cells were stressed for 20 min and then immunostained as previously described (Rogers and Rogers, 2008; Schoborg *et al*., 2013). In brief, cells were fixed with 4% PFA for 10 min at RT, rinsed 3x with PBS, permeabilized with 0.2% Triton X-100 for 5 min, and blocked with 3% nonfat milk for 10 min at room temperature. Primary antibodies were diluted in 3% nonfat milk, and coverslips were incubated for 1 hour at room temperature in a humidified chamber followed by a 3x wash with PBS/0.1% Triton X-100 for 10 min each. Secondary antibodies were then diluted in 3% nonfat milk and incubated for 1 hour at room temperature, and coverslips were washed as described. 0.5 µg/ml DAPI was added to counterstain DNA; slides were then rinsed twice with PBS, and mounted in Vectashield.

### Cytology for mitotic indexing

Mitotic spreads from larval brains were scored for mitotic indices as described in (Gatti and Goldberg, 1991). In brief, larval brains were dissected in insect media (HyClone SFX; Thermo Fisher Scientific), incubated in 0.5% sodium citrate, pH 6.0, for 10 minutes followed by fixation with 4% PFA; 50% acetic acid, and softly squashed between a coverslip and slide. Slides were stained with 4′, 6-diamidino-2-phenylindole (DAPI, 0.5 µg/ml) and were mounted in Vectashield mounting medium (Vector Laboratories).

### X-ray sensitivity assessment

Mitotic chromosomes from larval brains were observed for the presence of aberrations following X-ray treatment and recovery in a similar manner as (Gatti and Goldberg, 1991; Merigliano *et al*., 2017). In short, third instar larvae, Oregon-R and *hipp1^ΔCLD^*, were irradiated with 7.5 Gy. Irradiation was performed in a Rad-Source RS-2000 Biological Irradiator. 2 hours after X-ray exposure, larval brains were dissected and placed in insect media (HyClone SFX; Thermo Fisher Scientific) supplemented with 0.1 mM colchicine for 1 hour, followed by a 10 minute incubation in 0.5% sodium citrate, pH 6.0. Brains were fixed with 4% PFA; 50% acetic acid. After the incubations, brains were then softly squashed and stained with 4′, 6-diamidino-2-phenylindole (DAPI, 0.5 µg/ml), rinsed twice with PBS and mounted in Vectashield mounting medium (Vector Laboratories). Brains were mounted on individual slides and observed. Slides were scanned for metaphasic nuclei and approximately 50 metaphases were collected for each slide. Metaphases were then scored for the presence of CABs.

### EdU incorporation and detection

Salivary glands or brains dissected from larvae were labeled with EdU according to the manufacturer’s protocol (Click-iT EdU Alexa Fluor 488 Imaging Kit; C10337, Invitrogen). In brief, tissues were incubated in 10 µM EdU diluted in insect media (HyClone SFX; Thermo Fisher Scientific) for the indicated amount of time. The tissues were fixed in 4% PFA; 50% acetic acid and adhered to a microscope slide. Slides were treated with the Click-iT reaction cocktail containing the Alexa Fluor azide for 30 minutes. Slides were washed in a blocking solution (PBS, 0.1% NP40, 3% nonfat milk) and labeled with the indicated primary and secondary antibodies and DAPI (0.5 µg/ml) before observation.

### Microscopy

Slides were analyzed using a wide-field epifluorescence microscope (DM6000 B; Leica) equipped with a charge-coupled device camera (ORCA-ER; Hamamatsu Photonics) and an HCX Plan Apochromat (Leica) 40X or 100X/1.35 NA oil immersion objective. Image acquisition was performed using SimplePCI (v6.6; Hamamatsu Photonics). Image brightness and contrast adjustments were performed using Fiji (Schindelin et al., 2012). Samples were processed and imaged under identical conditions of immunostaining, microscope, camera, and software settings.

### BrdU labeling and analysis

For the BrdU incorporation assay, brains dissected from third instar larvae were incubated for 1 hour in 0.1 mg/ml BrdU at room temperature. Tissues were then fixed with 4% PFA; 50% acetic acid, washed in PBS, and incubated in blocking solution. Brains were then labeled with 2 μg/ml BrdU primary antibody (Developmental Studies Hybridoma Bank, G3G4) and DAPI (0.5 µg/ml). BrdU signal intensity was quantified using Image J analysis software. Regions of interest (ROIs) were determined for each nucleus using signal from the DAPI channel. Background subtraction using a rolling ball algorithm was performed prior to taking measurements. To sort cells based on DNA content, DAPI intensity was measured from ROIs determined by DAPI staining. Line plots were generated by quantifying the frequency of nuclei that fell within ranges of fluorescent intensity.

## Acknowledgements

We thank Judith Kassis for her generous Wapl antibody gift. BrdU antibodies were obtained from the Developmental Studies Hybridoma Bank, created by the NICHD of the NIH and maintained at The University of Iowa, Department of Biology, Iowa City, IA 52242. We also thank the Biochemistry and Cellular and Molecular Biology Department, the College of Arts and Sciences, and Office of Research at the University of Tennessee for support in addition to the US Public Health Service Awards GM78132-2 and MH108956 from the National Institutes of Health and from National Science Foundation (0616081) to M. Labrador.

